# Effects of source sample amount on biodiversity surveys of bacteria, fungi, and nematodes in soil ecosystems

**DOI:** 10.1101/2022.05.26.493541

**Authors:** Takuya Kageyama, Hirokazu Toju

## Abstract

Bacteria, fungi, and nematodes are major components of soil ecosystems, playing pivotal roles in belowground material cycles and biological community processes. A number of studies have recently uncovered the diversity and community structure of those organisms in various types of soil ecosystems based on DNA metabarcoding (amplicon sequencing). However, because each of the most previous studies examined only one or two of the three organismal groups, it remains an important challenge to reveal the entire picture of soil community structure. We herein examined how we could standardize DNA extraction protocols for simultaneous DNA metabarcoding of bacteria, fungi, and nematodes. Specifically, in an Illumina sequencing analysis of forest and farmland soil samples, we performed DNA extraction at five levels of soil-amount settings (0.5 g, 2 g, 5 g, 10 g, and 20 g). We then found that DNA extraction with the 0.5-g soil setting, which had been applied as default in many commercial DNA extraction kits, could lead to underestimation of *α*-diversity in nematode community. The detected family-level diversity of nematodes, in particular, increased with increasing soil amounts in DNA extraction. Our analyses also indicated that dissimilarity (*β*-diversity) of community structure among replicate samples were higher in fungi and nematodes than in bacteria, suggesting difference in basic levels of spatial heterogeneity among the three organismal groups. These findings provide platforms for optimizing DNA metabarcoding protocols commonly applicable to major groups of organisms in soil ecosystems.

## INTRODUCTION

Bacteria, fungi, and nematodes occupy a large amount of biomass in the soil, playing pivotal ecosystem functions (Fierer et al., 2007; Bardgett and van der Putten, 2014; Nielsen et al., 2015; Wall et al., 2015; Bahram et al., 2018; Bar-On et al., 2018; Flemming and Wuertz, 2019; van den Hoogen et al., 2019). In uncovering the tremendous diversity of those soil organisms, DNA metabarcoding (amplicon sequencing) based on high-throughput sequencing has been increasingly applied as a standard approach (Caporaso et al., 2012; Reuter et al., 2015). Through the recent enrichment of reference DNA databases of bacterial 16S rRNA, fungal internal transcribed spacer (ITS), and nematode 18S rRNA sequences, it has become possible to gain large datasets of soil biodiversity without longtime taxonomic expertise or experiences in microbial isolation (Quast et al., 2013a; Nilsson et al., 2019).

In expanding the use of DNA metabarcoding of soil biodiversity, it is of particular importance to develop standard protocols applicable to broad ranges of situations and organisms. For example, universal primers with high taxonomic coverage have been proposed for each of bacterial 16S rRNA (Klindworth et al., 2013; Thijs et al., 2017), fungal ITS (Schoch et al., 2012; Lindahl et al., 2013; op de Beeck et al., 2014), and nematode 18S rRNA regions (Sapkota and Nicolaisen, 2015; Ahmed et al., 2019; Waeyenberge et al., 2019; Sikder et al., 2020; Kenmotsu et al., 2021). For the bioinformatic processes, rapid and accurate algorithms for inferring operational taxonomic units (OTUs) or amplicon sequence variants (ASVs) have been implemented (Callahan et al., 2016). Likewise, theories on reference database search (Tanabe and Toju, 2013) and taxonomic assignment (Huson et al., 2007) have been proposed to allow automatic molecular identification. While all these processes are keys to reliability and reproducibility of DNA metabarcoding, results of molecular-based biodiversity inventories are considered to depend greatly on DNA extraction methods (Thompson et al., 2017; Zielińska et al., 2017; Francioli et al., 2021). In particular, the amount of the soil subjected to DNA extraction possibly have major impacts on the *α-* and β*-*diversity estimates of metabarcoding samples.

Because soil environments such as pH, humidity, and nutrient levels vary within ecosystems, the taxonomic compositions of soil organisms often show high spatial heterogeneity (Franklin and Mills, 2003; Fierer, 2017; Flemming and Wuertz, 2019). Therefore, DNA extraction from small amounts of soil samples may lead to the underestimation of biodiversity or low reproducibility in the reconstruction of community structure. To date, a number of studies have been conducted to evaluate how soil amounts in DNA extraction can affect DNA metabarcoding results of bacteria and/or fungi (Kang and Mills, 2006; Morise et al., 2012; Song et al., 2015; Penton et al., 2016). Meanwhile, few studies have examined the dependence of soil amounts on molecular analyses targeting nematode communities. Given the multicellularity and mobility of nematodes, their community structure is expected to be highly heterogeneous within soil ecosystems (Ferris et al., 1990; Ettema, 1998; Liu et al., 2019). Moreover, given the relatively large body size and low density of nematodes compared to those of bacteria, DNA extraction protocols optimized for bacteria may lead to the failure of nematode diversity profiling (Wiesel et al., 2015). Consequently, in developing DNA metabarcoding methods for simultaneous surveys of bacterial, fungal, and nematode communities in soil ecosystems, molecular experimental protocols need to be standardized by exploring optimal soil amounts in DNA extraction.

In this study, we examined how DNA metabarcoding data could be influenced by the amount of the soil subjected to DNA extraction. We collected soil specimens in forest and farmland (soybean field) ecosystems and then performed DNA extraction at five levels of soil-amount settings (0.5 g, 2 g, 5 g, 10 g, and 20 g). Based on the Illumina sequencing outputs, the number of detected taxa (e.g., genera and families) were compared among the five soil-amount classes for each ecosystem type (forest or farmland) for each of the bacterial 16S rRNA, fungal ITS, and nematode 18S rRNA regions. We also evaluated how the level of the among-replicate-sample heterogeneity of community structure could differ among bacterial, fungal, and nematode datasets, gaining insights into difference in the basic level of spatial heterogeneity among the three organismal groups. The results shown in this study will provide bases for designing DNA metabarcoding protocols commonly applicable to major taxa in soil ecosystems.

## MATERIALS AND METHODS

### Study site and soil sampling

Soil samples were collected on October 29, 2021, from a temperate secondary forest (34.972 ºN, 135.959 ºE) and an experimental soybean field (34.972 ºN, 135.958 ºE) of the Center for Ecological Research, Kyoto University. In the secondary forest, which was dominated by *Quercus serrata* (Fagaceae), litter on the soil surface was removed before sampling. Then, at each of the eight sampling positions set at 8-m intervals, three soil core (diameter = 3 cm, depth = 10 cm) samples collected within a 32 cm^2^ area were mixed in a plastic bag. In the soybean field, soil cores were collected at each of the eight sampling positions set at > 8-m intervals. The eight forest soil samples and the eight soybean-field soil samples were stored at -25 °C in the laboratory until DNA extraction.

### DNA extraction

Each soil sample was well mixed and subjected to the following DNA extraction processes with five alternative soil amount settings: specifically, DNA extraction from 0.5 g, 2 g, 5 g, 10 g, and 20 g of wet soil was conducted. For the 0.5-g setting, soil DNA was directly processed with Extrap Soil DNA Kit Plus Ver.2 (BioDynamics Laboratory Inc., Tokyo) based on the beads-beating protocol provided by the manufacturer. For the remaining four settings (2 g, 5 g, 10 g, and 20 g), each soil sample soaked in extraction buffer [10mM Tris-HCl (pH 8.5), 1mM EDTA, 1% SDS] was shaken with 4-mm, 1-mm, and 0.5-mm zirconium beads (Asone, Osaka) in a 50-mL centrifuge tube at 6.5 m/sec for 60 seconds using FastPrep 24 (MP Biomedicals, USA). After centrifugation, 500 µL of the supernatant was subjected to DNA extraction using Extrap Soil DNA Kit without the default beads-beating step. In total, 80 DNA template samples were obtained [2 ecosystem types (forest or soybean field) × 8 samples × 5 soil-amount classes].

### DNA amplification and sequencing

Profiling of biodiversity was performed by targeting bacteria, fungi, and nematodes. For the amplification of the 16S rRNA V4 region of bacteria, the set of the forward primer 515f (Caporaso et al., 2011) and the reverse primer 806rB (Apprill et al., 2015) were used. The primers were fused with 3–6-mer Ns for improved Illumina sequencing quality (Lundberg et al., 2013) and Illumina sequencing primers as detailed in a previous study (Toju et al., 2019). PCR was performed at a 10-µL-scale protocol of KOD ONE PCR Master Mix (TOYOBO, Osaka) with the temperature profile of 35 cycles at 98 ºC for 10 seconds (denaturation), 60 ºC for 5 seconds (annealing of primers), and 68 ºC for 5 seconds (extension), and a final extension at 68 ºC for 2 minutes. The ramp rate through the thermal cycles was set to 1 ºC/sec to prevent the generation of chimeric sequences (Stevens et al., 2013).

Likewise, the ITS1 region of fungi was amplified using the set of the forward primer ITS1F_KYO1 and the reverse primer ITS2_KYO2 (Toju et al., 2012). PCR was performed using the Illumina-sequencing fusion primer design mentioned above at a 10-µL-scale protocol of KOD ONE PCR Master Mix (TOYOBO, Osaka) with the temperature profile of 35 cycles at 98 ºC for 10 seconds, 52 ºC for 5 s seconds, and 68 ºC for 5 seconds, and a final extension at 68 ºC for 2 minutes (ramp rate = 1 ºC/sec).

For the amplification of nematodes, we redesigned the primer sets targeting nuclear 18S rRNA region (Kenmotsu et al., 2021) to improve taxonomic coverage. Specifically, using the aligned 18S rRNA sequences of major nematode taxa [downloaded from the NCBI database (https://www.ncbi.nlm.nih.gov/) on November 8, 2020; Supplementary Data S1], we designed the new forward primer NF1_rv (5’-GGT GCA TGG CCG TTC TTA GTT -3’) and reverse primer Nem18SV8_rv (5’-GTG TGT ACA AAK GGC AGG GAC -3’). The PCR for Illumina sequencing was performed with the same thermal cycle protocol used in the analysis of the fungal ITS1 region.

The PCR products of the bacterial 16S rRNA, fungal ITS1, and nematode 18S rRNA regions were respectively subjected to the additional PCR step for linking Illumina sequencing adaptors and 8-mer sample identifier indexes (Hamady et al., 2008) with the amplicons as detailed elsewhere (Toju et al., 2019). The temperature profile in the PCR was 7 cycles at 98 ºC for 10 seconds, 55 ºC for 5 seconds, and 68 ºC for 5 seconds, and a final extension at 68 ºC for 2 minutes. The PCR products of the 80 samples were then pooled for each of the 16S rRNA, fungal ITS1, and nematode 18S rRNA regions after a purification/equalization process with the AMPureXP Kit (Beckman Coulter, Inc., Brea). Primer dimers, which were shorter than 200 bp, were removed from the pooled library by supplemental purification with AMpureXP: the ratio of AMPureXP reagent to the pooled library was set to 0.8 (v/v) in this process. The sequencing libraries of the three regions were processed in an Illumina MiSeq sequencer (15% PhiX spike-in). Because the quality of forward sequences is generally higher than that of reverse sequences in Illumina sequencing, we optimized the MiSeq run setting in order to use only forward sequences. Specifically, the run length was set 271 forward (R1) and 31 reverse (R4) cycles to enhance forward sequencing data: the reverse sequences were used only for discriminating between bacterial 16S, fungal ITS1, and nematode 18S rRNA sequences based on the sequences of primer positions.

### Bioinformatics

In total, 13,039,352 sequencing reads were obtained in the Illumina sequencing. The raw sequencing data were converted into FASTQ files using the program bcl2fastq 1.8.4 distributed by Illumina. For each of the 16S rRNA, fungal ITS1, and nematode 18S rRNA regions, the output FASTQ files were demultiplexed using Claident v0.9.2022.01.26 (Hamady et al., 2008; Tanabe and Toju, 2013). The removal of low-quality sequences and ASV inferences were done using DADA2 (Callahan et al., 2016) v1.17.5 of R v3.6.3 (R Core Team, 2021). The mean number of filtered sequencing reads obtained per sample was 2,758, 1,851, and 2,113 for the bacterial, fungal, and nematode datasets, respectively. Taxonomic annotation of bateria and fungi was conducted based on the SILVA 138 SSU (Quast et al., 2013) and UNITE version 8.2 (Tedersoo et al., 2018) the assignTaxonomy function of DADA2. For the taxonomic annotation of nematodes, the five-nearest-neighbor method (Tanabe and Toju, 2013) was applied to the NCBI nucleotide sequence database bundled with Claident v0.9.2022.01.26. ASVs that were not assigned to the domain Bacteria, the kingdom Fungi, and the phylum Nematoda were removed from the 16S rRNA, ITS1, and 18S rRNA datasets, respectively. For each target organismal group (Bacteria, Fungi, and Nematoda), we then obtained a sample × ASV matrix, in which a cell entry depicted the number of sequencing reads of an ASV in a sample (Supplementary Data S2). The samples with less than 1,000 reads were discarded from the matrices. The sequencing data were deposited to DNA Data Bank of Japan (DDBJ) (accession no.: DRA014170).

### Statistical analysis

To examine how ASV richness increased with increasing number of sequencing reads, the iNEXT v.2.0.20 package of R for drawing the interpolation and extrapolation curves was used (Hsieh et al., 2016). The Hill numbers of bacterial, fungal, and nematode ASVs were then calculated for each soil-amount class of each soil sample. Based on the Hill number data, the *α*-diversity (family richness, Shannon diversity, and Simpson diversity) of bacteria, fungi, and nematodes were respectively examined in ANOVA models including ecosystem type (forest or soybean field) and soil-amount classes as explanatory variables. Multiple comparison was then performed with the Kruskal-Wallis test. The analysis was conducted for the community datasets at the ASV, genus, and family levels.

For each of the bacterial, fungal, and nematode datasets, the genus-, family-, and order-level taxonomic compositions of each sample was visualized as bar graphs after performing the rarefaction of the data using the vegan v.2.5.7 package of R (1,000 reads per sample). To compare the levels of spatial heterogeneity of community structure among bacteria, fungi, and nematodes, we calculated the β-diversity (Bray-Curtis dissimilarity) of family-level taxonomic compositions among replicate samples at each soil amount class. At each taxonomic level (genus or family), we constructed an ANOVA including among-sample β-diversity as a response variable and ecosystem type (forest or soybean field), organismic group (bacteria, fungi, or nematodes), and soil-amount class as explanatory variables.

## RESULTS

In total, 2,402 bacterial, 1,044 fungal, and 288 nematode ASVs were detected. The number of bacterial, fungal, and nematode genera were 234, 219, and 47, respectively. Likewise, the number of bacterial, fungal, and nematode families were 172, 157, and 38, respectively. For all the three organismic groups, interpolation and extrapolation curve were saturated: sample coverage was higher than 0.9 in all the analyzed samples (Figure 1). The taxonomic compositions of the samples were visualized in Figures 2-4 and Supplementary Figures S1–6).

**FIGURE 1.**
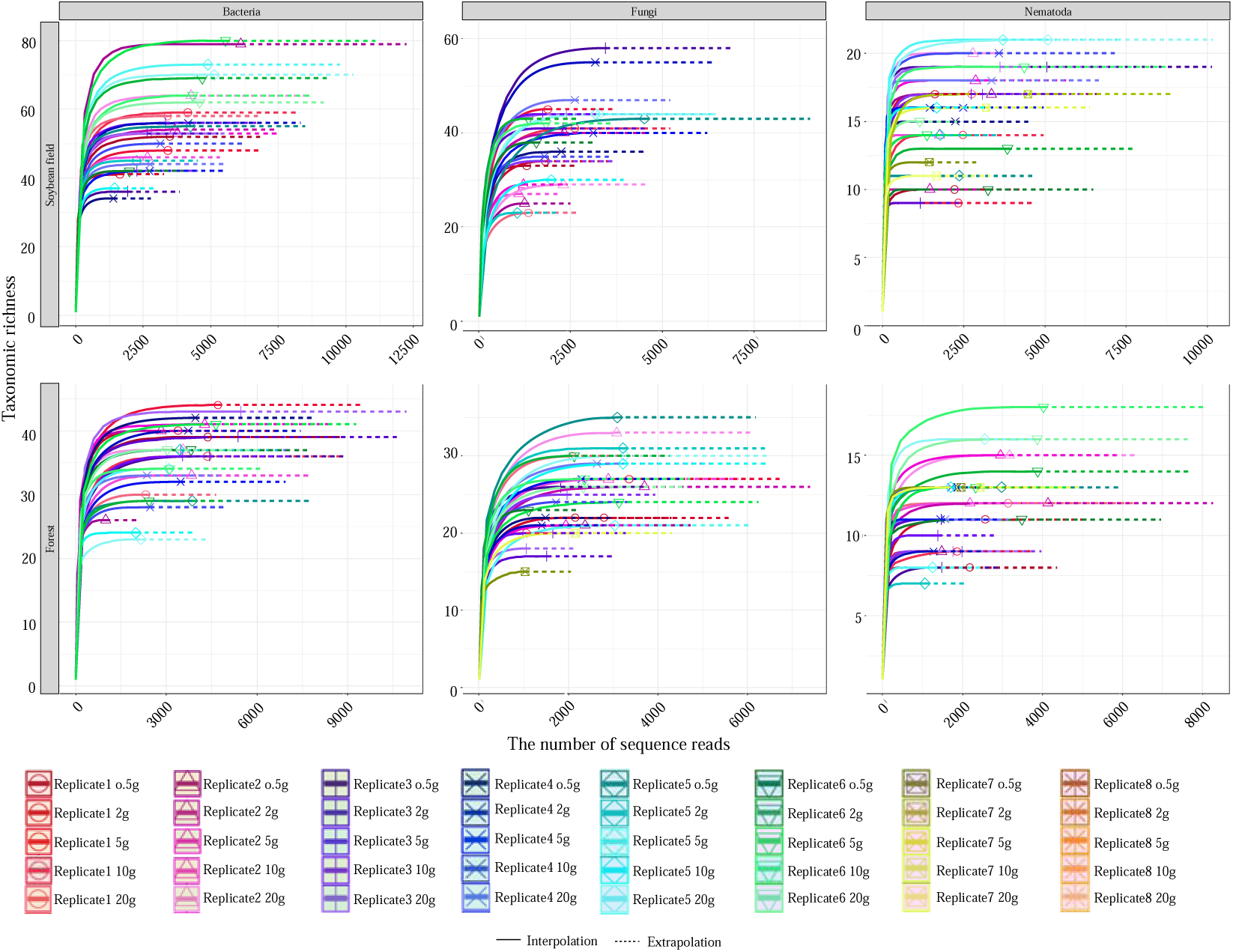
Interpolation and extrapolation curves of bacteria, fungi, and nematodes. The relationships between the number of sequencing reads and the number of detected families are shown for the bacterial 16S rRNA, fungal ITS, and nematode 18S rRNA datasets for each ecosystem type (forest or soybean field). Interpolation and extrapolation curves are shown in solid and dashed lines, respectively.

**FIGURE 2.**
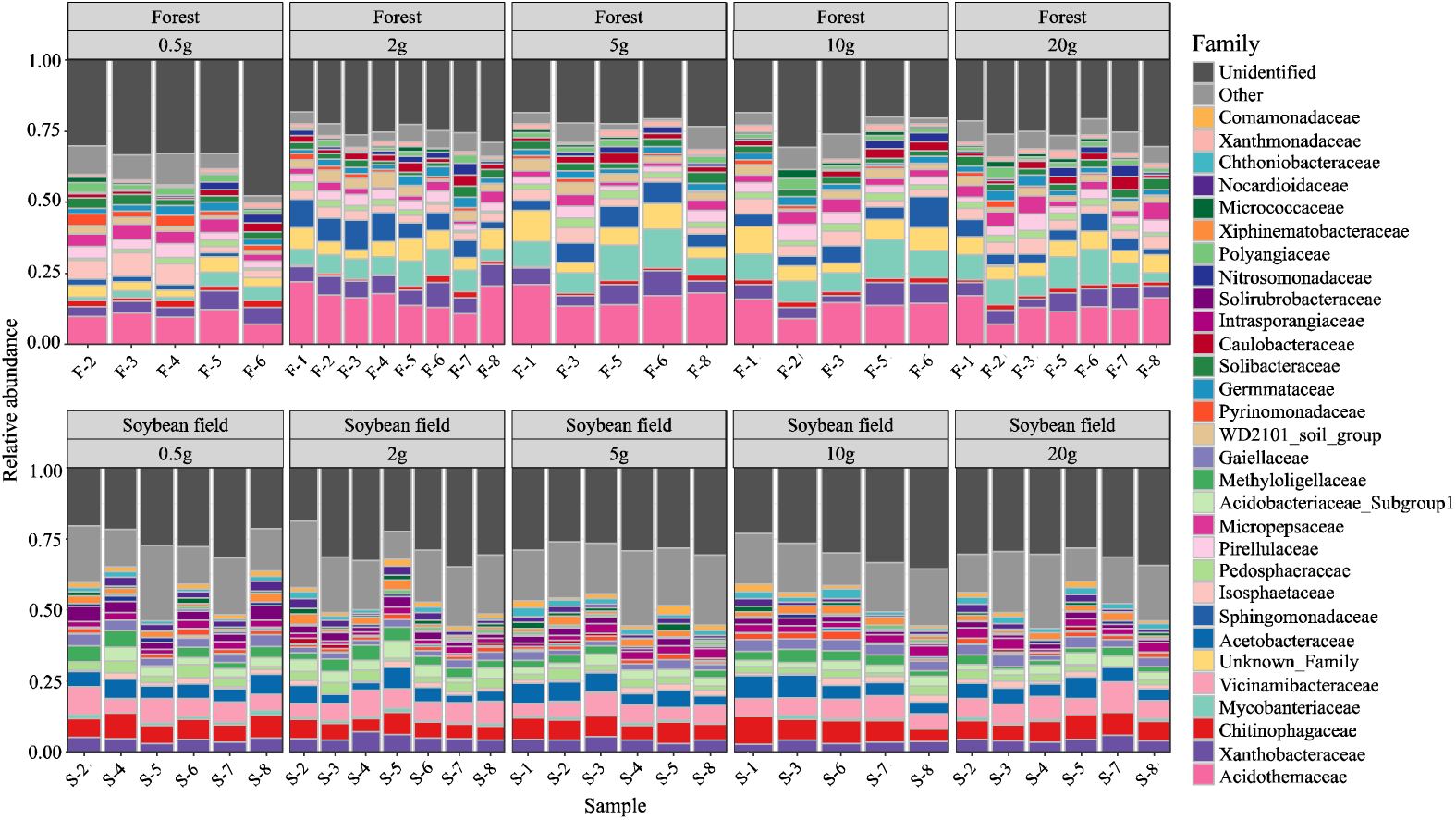
Family-level taxonomic compositions of bacteria. For each replicate sample of each ecosystem type (forest or soybean field), family-level compositions of the sequencing data are shown at each soil-amount class. Samples with less than 2000 sequencing reads were omitted. See Supplementary Figures S1 and S4 for genus- and order-level taxonomic compositions.

**FIGURE 3.**
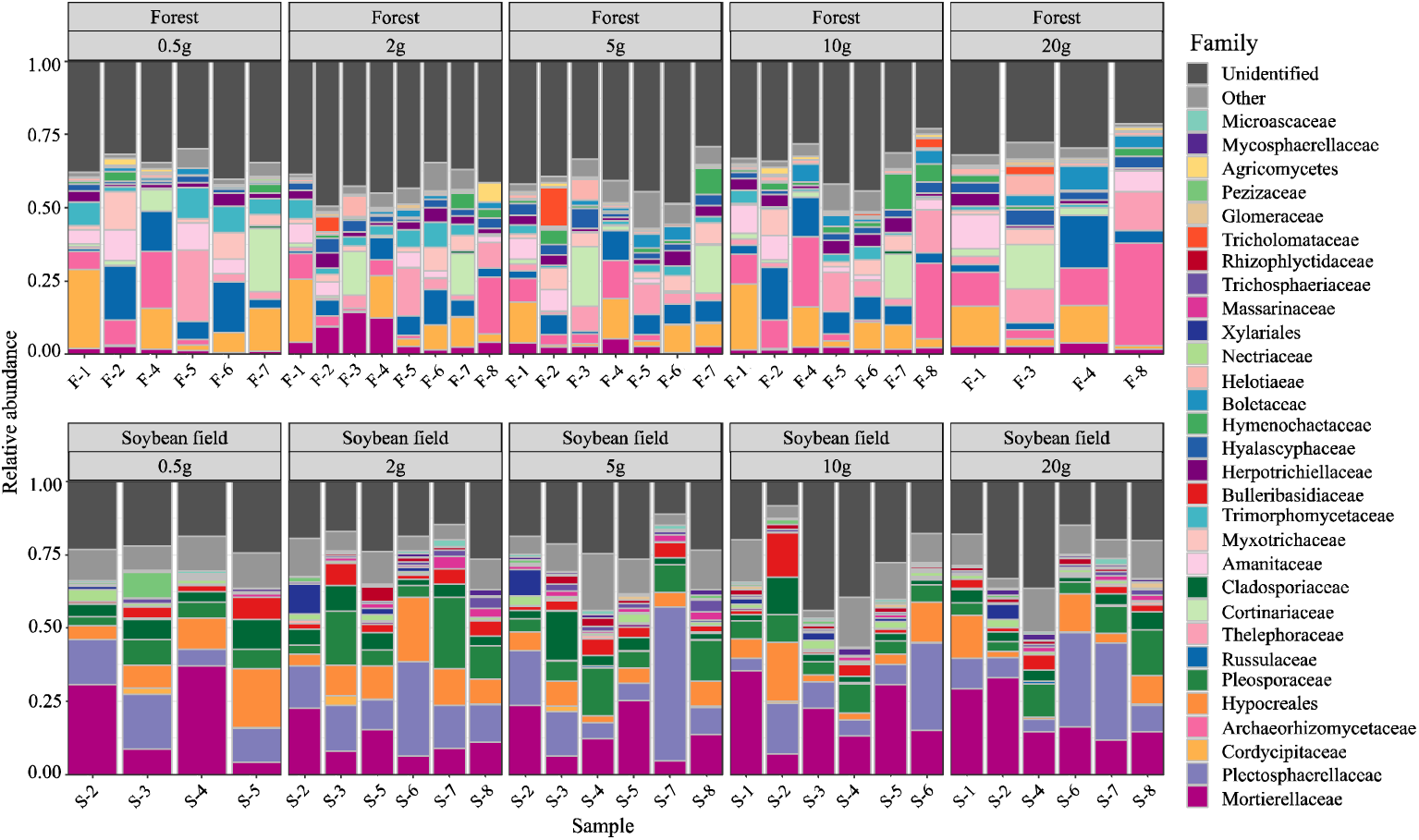
Family-level taxonomic compositions of fungi. For each replicate sample of each ecosystem type (forest or soybean field), family-level compositions of the sequencing data are shown at each soil-amount class. Samples with less than 2000 sequencing reads were omitted. See Supplementary Figures S2 and S5 for genus- and order-level taxonomic compositions.

**FIGURE 4.**
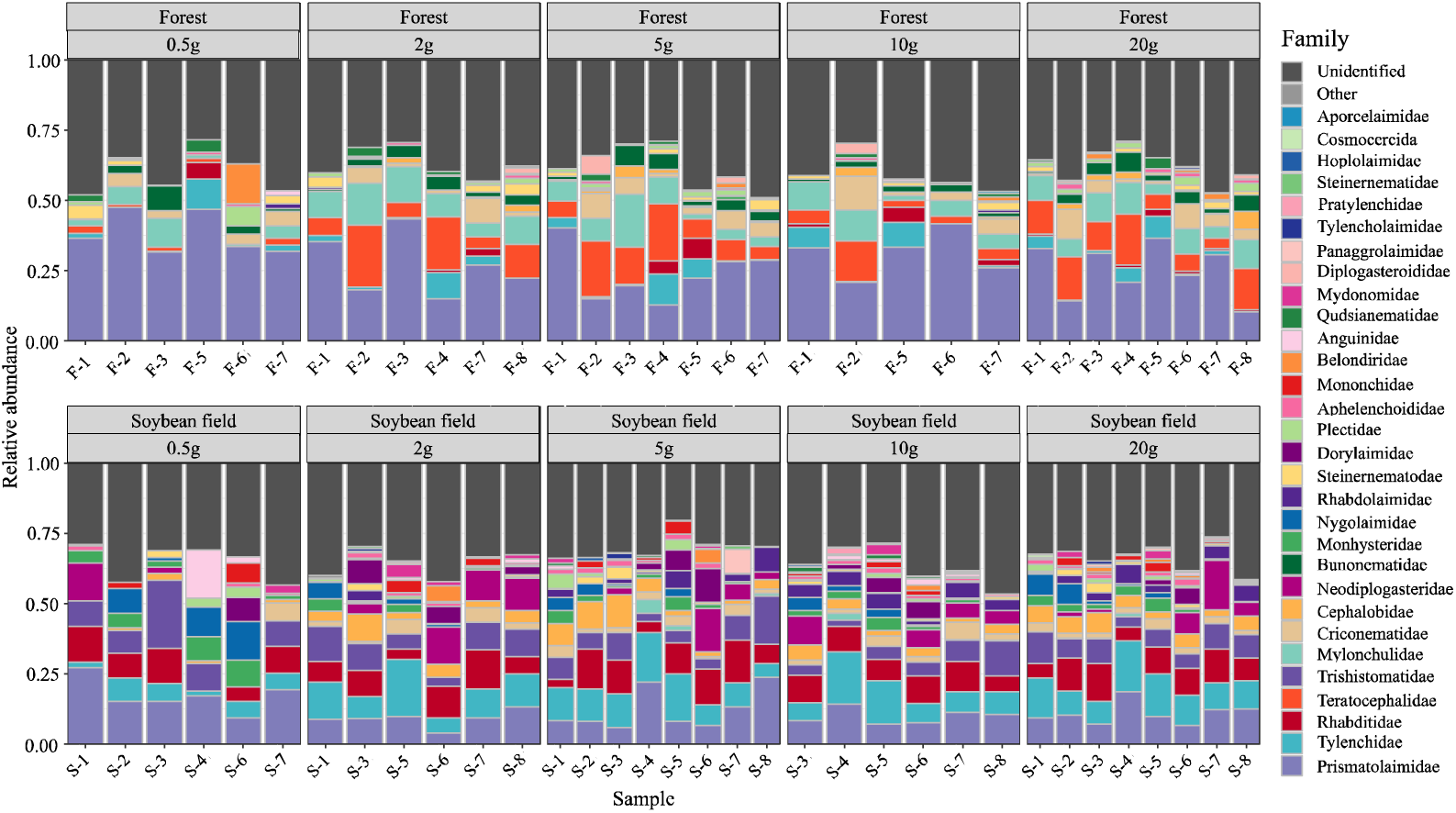
Family-level taxonomic compositions of nematodes. For each replicate sample of each ecosystem type (forest or soybean field), family-level compositions of the sequencing data are shown at each soil-amount class. Samples with less than 2000 sequencing reads were omitted. See Supplementary Figures S3 and S6 for genus- and order-level taxonomic compositions.

For bacteria and fungi, soil amount in DNA extraction did not have significant effects on the *α*-diversity estimates at the family level, while the interaction between ecosystem type (forest or soybean field) and soil amount had significant effects on the Shannon and Simpson diversity of bacteria (Table 1).

**TABLE 1.**
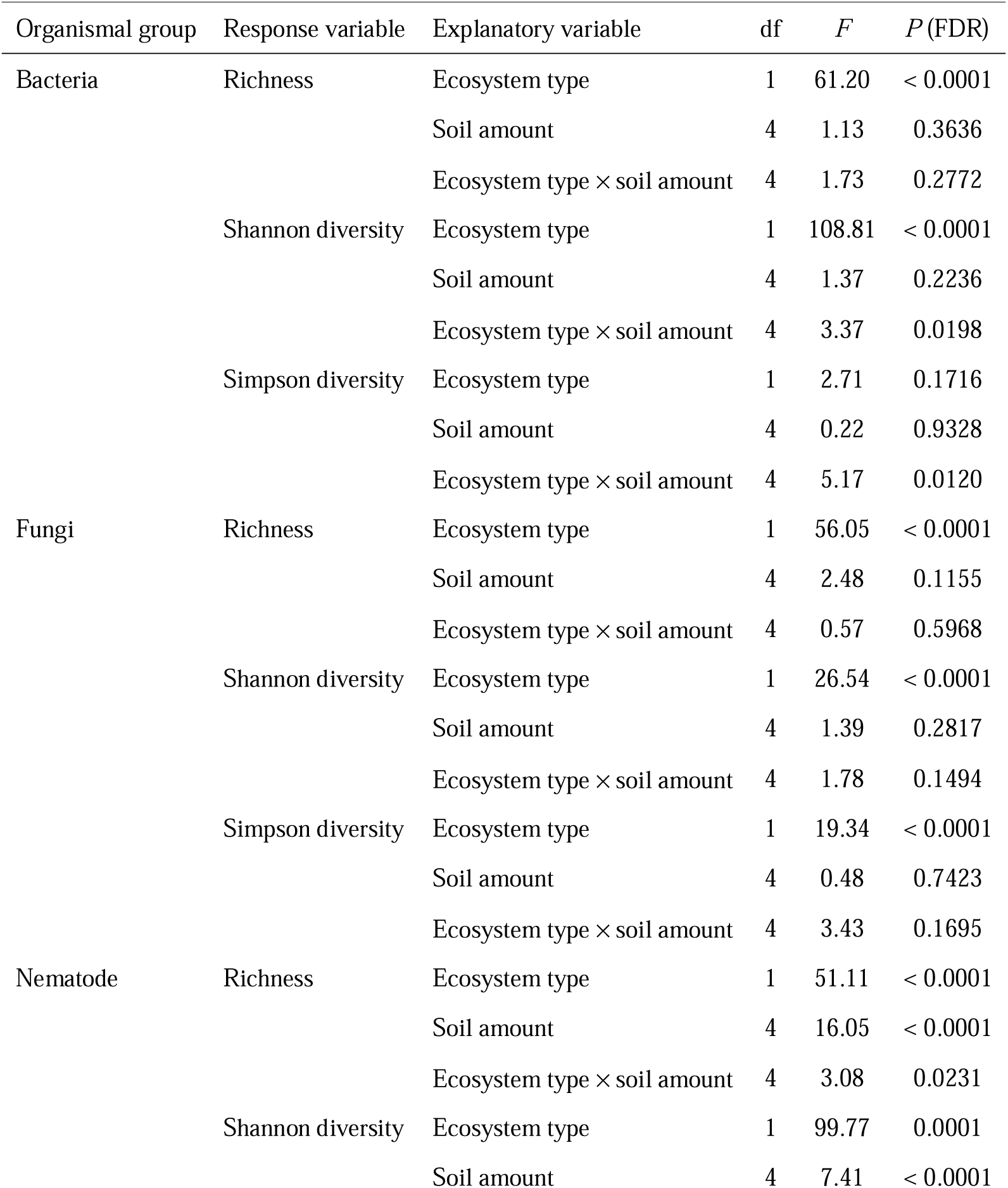
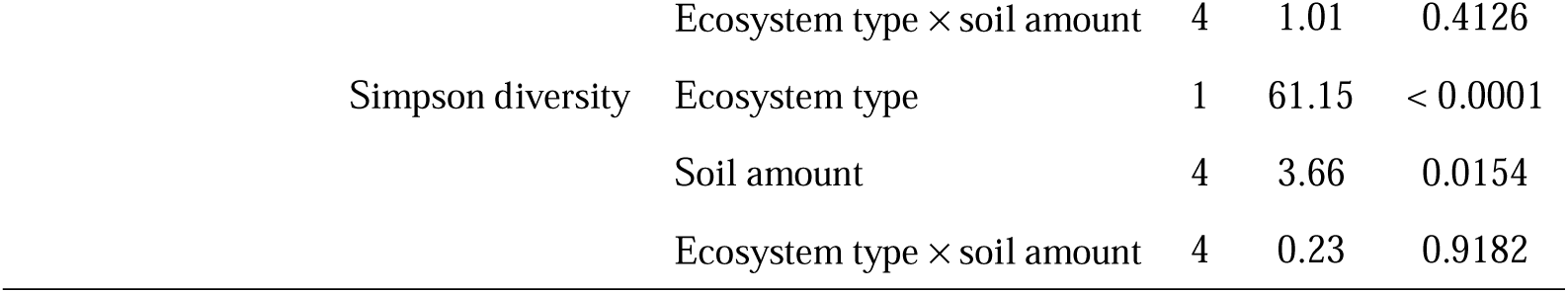
ANOVA of *α*-diversity. For each organismal group (bacteria, fungi, or nematodes), an ANOVA model of family-level *α*-diversity was constructed based on the number of families (richness), Shannon diversity, or Simpson diversity. The explanatory variables included win each model were ecosystem type (forest or soybean field), soil amount (2 g, 5 g, 10 g, and 20 g), and interaction term of the two factors. False discovery rates (FDR) were calculated for each organismal group.

In the nematode dataset, significant effects of soil amount on taxonomic richness, Shannon diversity, and Simpson diversity were observed (Table 1; Figure 5). Moreover, all the *α*-diversity indices had higher values at 0.5-g than at 20-g DNA extraction settings for the soybean field dataset in terms of Kruskal-Wallis test (Figure 5). The same contrast between 0.5-g and 20-g settings was observed in the analysis of taxonomic richness and Shannon diversity for the forest dataset (Figure 5). Qualitatively similar results were obtained at the ASV and genus levels (Supplementary Figure S7).

**FIGURE 5.**
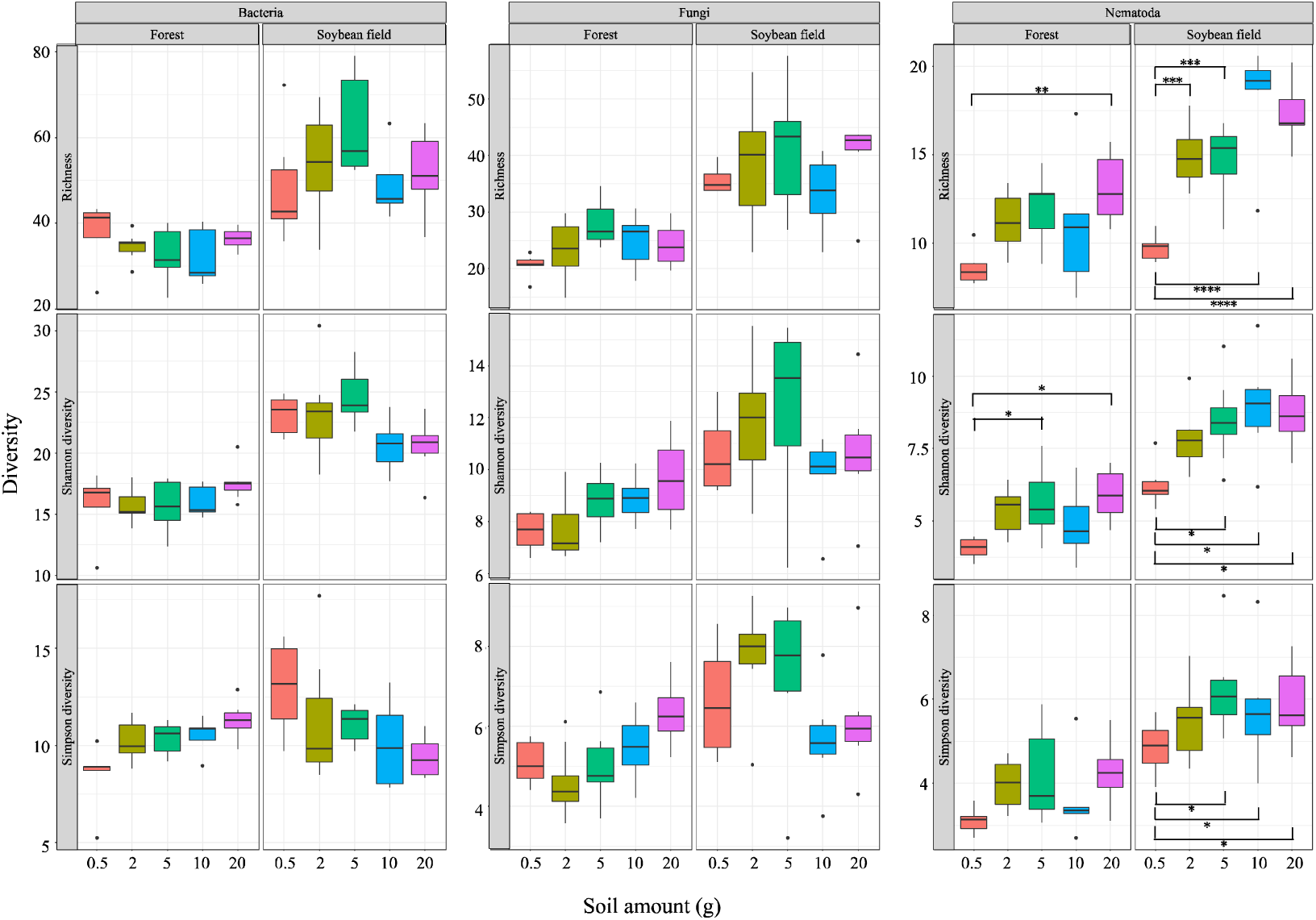
Effects of soil amount in DNA extraction on family-level *α*-diversity. For each of the bacterial 16S rRNA, fungal ITS, and nematode 18S rRNA datasets, taxonomic richness, Shannon diversity, and Simpson diversity at the family level are shown with boxplots for ecosystem type (forest or soybean field). See Table 1 for ANOVA results. For each ecosystem type, statistically significant difference inferred by Kruskal-Wallis test is indicated by asterisks. See Supplementary Figure S7 for ASV- and genus-level results.

The spatial heterogeneity (dissimilarity) in taxonomic compositions among replicate samples significantly varied depending on ecosystem type, soil amount in DNA extraction, and organismal groups as well as on the interaction terms of them (Table 2). Among the examined factors, organismal groups (bacteria, fungi, or nematodes) had the strongest effects (the largest *F* value) on among-sample community dissimilarity. Specifically, the spatial heterogeneity of taxonomic compositions was the highest for fungi and the lowest for bacteria (Figure 6). The nematode dataset displayed intermediate levels of spatial heterogeneity in both the forest and soybean field ecosystems (Figure 6).

**TABLE 2.** ANOVA of spatial heterogeneity in community structure among replicate samples. An ANOVA model of Bray-Curtis dissimilarity of family-level taxonomic compositions among replicate samples was constructed. The explanatory variables included were ecosystem type (forest or soybean field), organismal group (bacteria, fungi, or nematodes), soil amount (2 g, 5 g, 10 g, and 20 g), and interaction terms of the three factors.

| **Explanatory variable** | **df** | ***F*** | ***P* (FDR)** |
| --- | --- | --- | --- |
| **Ecosystem type** | 1 | 45.55 | < 0.0001 |
| **Organismal group** | 2 | 243.19 | < 0.0001 |
| **Soil amount** | 4 | 15.87 | < 0.0001 |
| **Ecosystem type × organism group** | 2 | 8.50 | 0.0459 |
| **Organism group × soil amount** | 8 | 4.53 | < 0.0001 |
| **Soil amount × ecosystem type** | 4 | 2.44 | 0.0002 |
| **Ecosystem type × organism group × soil amount** | 8 | 2.39 | 0.0157 |

**FIGURE 6.**
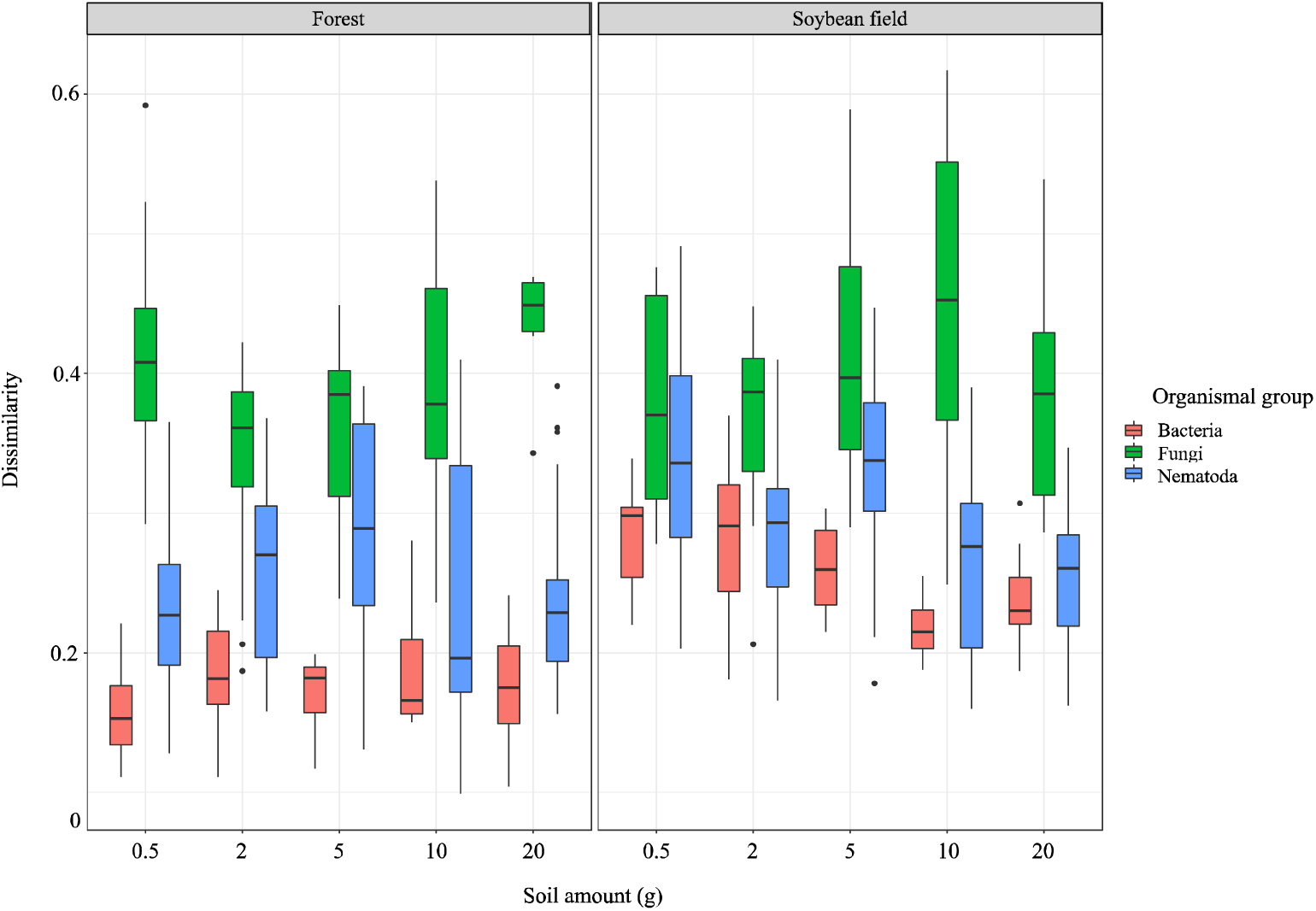
Dissimilarity of family-level taxonomic compositions among replicate samples. For each ecosystem type (forest or soybean field), Bray-Curtis *β*-diversity of family-level community structure among replicate samples is compared among the bacterial, fungal, and nematode datasets. See Table 2 for ANOVA results.

## DISCUSSION

We here examined how the amount of soil subjected to DNA extraction affects the results of biodiversity surveys targeting three major groups of soil organisms, namely, bacteria, fungi, and nematodes. We then found that the minimal amount of soil required for DNA metabarcoding could vary among these three organismal groups as detailed bellow.

For bacteria and fungi, there have been several studies evaluating potential effects of soil sample amount on DNA metabarcoding-based biodiversity inventories. Among those studies, some suggested that 10 g of soil were ideally required to gain reproducible ASV richness data (Kang and Mills, 2006; Penton et al., 2016), while others discussed smaller amounts of soil were enough for surveys of bacteria (and fungi) (Ranjard et al., 2003). In our study, the amount of soil subjected to DNA extraction did not directly affect the *α*-diversity estimates (i.e., taxonomic richness, Shannon diversity, and Simpson diversity) of bacteria and fungi (Table 1, Figure 5). Therefore, we conclude that DNA extraction from 0.5 g of soil and that from larger amounts of soil can yield qualitatively similar metabarcoding results in the analyses targeting bacteria and fungi. Meanwhile, taking into account the previous studies reporting potential bias related to DNA extraction from small amounts of soil (Kang and Mills, 2006; Penton et al., 2016), careful management of experimental protocols would be required in DNA extraction from 0.5. of soil.

While there have been a number of methodological studies on bacterial and fungal DNA metabarcoding, few studies have been conducted to examine the effects of soil sample amount on nematode diversity surveys. In research on soil nematode communities, the traditional approach is to isolate nematodes from the soil matrix by means of the Behrman method, elutriator and sucrose centrifugation, and filter separation, followed by microscopic morphological observations (Neher et al., 1995; Yeates et al., 1999; Ferris et al., 2012). This approach is usually taken based on sampling of several hundred grams of soil. Meanwhile, in recent years, DNA-based techniques have been recognized as an alternative approach and nematode community inventories from smaller amount of soil have started to be examined (Wiesel et al., 2015; Treonis et al., 2018; Quist et al., 2019; Zhang et al., 2021).

Our results indicated that soil amount in DNA extraction could considerably affect DNA metabarcoding results of nematodes (Figures 4 and 5). All the family-level *α*-diversity estimates increased with increasing soil amount, reaching maximum values at 20 g (Figure 5). Such increases in *α*-diversity with increasing soil amount in DNA extraction were observed as well at the ASV and genus levels (Supplementary Figure S7). These results indicate that biodiversity surveys of nematodes require higher amounts of soil than those of bacteria and fungi. This finding is as expected because nematodes [typically, from 0.1 to 2.5 mm long (Hodda, 2022)] are much larger than bacterial cells and fungal hyphae: thus, their population density in the soil may be much lower than microbes. Even if “environmental DNA” deriving from feces or remains of nematodes are detectable simultaneously with DNA from living nematode bodies, the scale of soil samples required for diversity inventories would be larger in nematodes than in bacteria and fungi (Peham et al., 2017). Given our data, we would suggest mixing as much soil as possible and then introducing subsampled 20 g of soil to lysis buffers for DNA extraction. Ultimately, the use of more soil in the lysis step would be ideal. However, 20 g may be the upper limit of molecular biological experiments given the size of centrifugeable tubes (≦ 50 mL). Consequently, setting replicate sampling points within each study site is of particular importance in nematode DNA barcoding even if DNA extraction is performed at the 20-g soil scale.

The results of the DNA metabarcoding conducted in this study also indicated that the level of dissimilarity (β-diversity) among replicate samples at each study site (forest or soybean field) varied among bacteria, fungi, and nematodes (Figure 6; Table 2). Specifically, as seen in the relatively homogeneous taxonomic compositions of bacteria among replicate samples (Figure 2), among-sample dissimilarity was lower in bacteria than in fungi and nematodes (Figure 6). In contrast, the family-level taxonomic compositions of fungi varied considerably among replicate samples, especially in the forest ecosystem (Figure 3), resulting in the highest levels of among-sample β-diversity (Figure 6). The intermediate levels of among-sample heterogeneity were observed for nematode taxonomic compositions (Figures 4 and 6). These results suggest that spatial ecological processes and microhabitat distributions of fungi and nematodes are more dynamic than those of bacteria in soil ecosystems. Consequently, for biodiversity inventories of multiple organismal groups, the number of replicate samples as well as intervals between sampling positions need to be optimized given the scale of spatial autocorrelations of community compositions (Štursová et al., 2016).

Although the above results are informative, it should be acknowledged that the optimal amount of soil for DNA metabarcoding potentially depend on soil and ecosystem types. For example, the level of spatial heterogeneity in fungal community structure can vary between grassland ecosystems and farmlands with continuous anthropogenic perturbations (e.g., hyphal disconnection by tillage). In addition, it has been reported that soil nematode density varies across continents depending on soil edaphic factors (van den Hoogen et al., 2019). Thus, for processing soil samples with highly heterogeneous community structure or low organismal densities, mechanical approaches for systematically mixing large amounts of soil before DNA extraction need to be developed in future studies.

Overall, we herein found that the standard DNA metabarcoding protocols developed for bacteria are not directly applicable to multicellular organisms in terms of the soil amount subjected to DNA extraction. Given that DNA-based inventories of biodiversity have become major tools in ecology, a next important step may be integration and comparison of diversity patterns among different groups of organisms. Towards the thorough understanding of metacommunity-scale dynamics and macroecological patterns of soil biodiversity, further attempts of technical standardization are necessary to target whole biomes including not only bacteria, fungi, and nematodes but also other invertebrates and protists.

## AUTHOR CONTRIBUTIONS

TK and HT designed the work. TK carried out the experiments. TK analyzed the data. TK and HT wrote the manuscript.

## ACKNOWLEDGEMENTS

This work was financially supported by JSPS Grant-in-Aid for Scientific Research (18H04009), JST FOREST (JPMJFR2048), and NEDO Moonshot Research and Development Program (JPNP18016) to HT.

## SUPPLEEMENTARY MATERIAL

The Supplementary Material for this article can be found online at [URL to be assigned by the publisher].

## REFERENCES

Ahmed, M., Back, M. A., Prior, T., Karssen, G., Lawson, R., Adams, I., et al. (2019). Metabarcoding of soil nematodes: the importance of taxonomic coverage and availability of reference sequences in choosing suitable marker(s). Metabarcoding and Metagenomics 3, 37–99. doi: 10.3897/mbmg.3.36408.

Apprill, A., Mcnally, S., Parsons, R., and Weber, L. (2015). Minor revision to V4 region SSU rRNA 806R gene primer greatly increases detection of SAR11 bacterioplankton. Aqua. Microb. Ecol. 75, 129–137. doi: 10.3354/ame01753.

Bahram, M., Hildebrand, F., Forslund, S. K., Anderson, J. L., Soudzilovskaia, N. A., Bodegom, P. M., et al. (2018). Structure and function of the global topsoil microbiome. Nature 560, 233–237. doi: 10.1038/s41586-018-0386-6.

Bardgett, R. D., and van der Putten, W. H. (2014). Belowground biodiversity and ecosystem functioning. Nature 515, 505–511. doi: 10.1038/nature13855.

Bar-On, Y. M., Phillips, R., and Milo, R. (2018). The biomass distribution on Earth. Proc Natl Acad Sci USA 115, 6506–6511. doi: 10.1073/pnas.1711842115.

Callahan, B. J., McMurdie, P. J., Rosen, M. J., Han, A. W., Johnson, A. J. A., and Holmes, S. P. (2016). DADA2: High-resolution sample inference from Illumina amplicon data. Nat. Methods 13, 581–583. doi: 10.1038/nmeth.3869.

Caporaso, J. G., Lauber, C. L., Walters, W. A., Berg-Lyons, D., Huntley, J., Fierer, N., et al. (2012). Ultra-high-throughput microbial community analysis on the Illumina HiSeq and MiSeq platforms. ISME J. 6, 1621–1624. doi: 10.1038/ismej.2012.8.

Caporaso, J. G., Lauber, C. L., Walters, W. A., Berg-Lyons, D., Lozupone, C. A., Turnbaugh, P. J., et al. (2011). Global patterns of 16S rRNA diversity at a depth of millions of sequences per sample. Proc Natl Acad Sci USA 108, 4516–4522. doi: 10.1073/pnas.1000080107.

Ettema, C. H. (1998). Soil nematode diversity: species coexistence and ecosystem function. J. Nematol. 30, 159–69. http://www.ncbi.nlm.nih.gov/pubmed/19274206.

Ferris, H., Mullens, T. A., and Foord, K. E. (1990). Stability and characteristics of spatial description parameters for nematode populations. J. Nematol. 22, 427–39. http://www.ncbi.nlm.nih.gov/pubmed/19287742.

Ferris, H., Sánchez-Moreno, S., and Brennan, E. B. (2012). Structure, functions and interguild relationships of the soil nematode assemblage in organic vegetable production. Appl. Soil Ecol. 61, 16–25. doi: 10.1016/j.apsoil.2012.04.006.

Fierer, N. (2017). Embracing the unknown: Disentangling the complexities of the soil microbiome. Nat. Rev. Micro. 15, 579–590. doi: 10.1038/nrmicro.2017.87.

Fierer, N., Bradford, M. A., and Jackson, R. B. (2007). Toward an ecologica classification of soil bacteria. Ecology 88, 1354–1364. doi: 10.1890/05-1839.

Flemming, H. C., and Wuertz, S. (2019). Bacteria and archaea on Earth and their abundance in biofilms. Nat. Rev. Micro. 17, 247–260. doi: 10.1038/s41579-019-0158-9.

Francioli, D., Lentendu, G., Lewin, S., and Kolb, S. (2021). DNA metabarcoding for the characterization of terrestrial microbiota—pitfalls and solutions. Microorganisms 9, 1–29. doi: 10.3390/microorganisms9020361.

Franklin, R. B., and Mills, A. L. (2003). Multi-scale variation in spatial heterogeneity for microbial community structure in an eastern Virginia agricultural field. FEMS Microbiol. Ecol. 44, 335–346. doi: 10.1016/S0168-6496(03)00074-6.

Hamady, M., Walker, J. J., Harris, J. K., Gold, N. J., and Knight, R. (2008). Error-correcting barcoded primers for pyrosequencing hundreds of samples in multiplex. Nat. Methods 5, 235–237. doi: 10.1038/nmeth.1184.

Hodda, M. (2022). Phylum Nematoda: a classification, catalogue and index of valid genera, with a census of valid species. Zootaxa 5114, 1–289. doi: 10.11646/zootaxa.5114.1.1.

Hsieh, T. C., Ma, K. H., and Chao, A. (2016). iNEXT: an R package for rarefaction and extrapolation of species diversity (Hill numbers). Methods Ecol. Evol. 7, 1451–1456. doi: 10.1111/2041-210X.12613.

Huson, D. H., Auch, A. F., Qi, J., and Schuster, S. C. (2007). MEGAN analysis of metagenomic data. Genome Res. 17, 377–386. doi: 10.1101/gr.5969107.

Kang, S., and Mills, A. L. (2006). The effect of sample size in studies of soil microbial community structure. J. Microbiol. Methods 66, 242–250. doi: 10.1016/j.mimet.2005.11.013.

Kenmotsu, H., Ishikawa, M., Nitta, T., Hirose, Y., and Eki, T. (2021). Distinct community structures of soil nematodes from three ecologically different sites revealed by high-throughput amplicon sequencing of four 18S ribosomal RNA gene regions. PLoS ONE 16. doi: 10.1371/journal.pone.0249571.

Klindworth, A., Pruesse, E., Schweer, T., Peplies, J., Quast, C., Horn, M., et al. (2013). Evaluation of general 16S ribosomal RNA gene PCR primers for classical and next-generation sequencing-based diversity studies. Nucleic Acids Res. 41. doi: 10.1093/nar/gks808.

Lindahl, B. D., Nilsson, R. H., Tedersoo, L., Abarenkov, K., Carlsen, T., Kjøller, R., et al. (2013). Fungal community analysis by high-throughput sequencing of amplified markes - a user’s guide. New Phytol. 199, 288–299. doi: 10.1111/nph.12243.

Liu, T., Hu, F., and Li, H. (2019). Spatial ecology of soil nematodes: Perspectives from global to micro scales. Soil Biol. Biochem. 137. doi: 10.1016/j.soilbio.2019.107565.

Lundberg, D. S., Yourstone, S., Mieczkowski, P., Jones, C. D., and Dangl, J. L. (2013). Practical innovations for high-throughput amplicon sequencing. Nat. Methods 10, 999–1002. doi: 10.1038/nmeth.2634.

Morise, H., Miyazaki, E., Yoshimitsu, S., and Eki, T. (2012). Profiling Nematode Communities in Unmanaged Flowerbed and Agricultural Field Soils in Japan by DNA Barcode Sequencing. PLoS ONE 7. doi: 10.1371/journal.pone.0051785.

Neher, D. A., Peck, S. L., Rawlings, J. O., and Campbell, C. L. (1995). Measures of nematode community structure and sources of variability among and within agricultural fields. Plant and Soil 170, 167–181. doi: 10.1007/BF02183065.

Nielsen, U. N., Wall, D. H., and Six, J. (2015). Soil Biodiversity and the Environment. Annu. Rev. Environ Resour. 40, 63–90. doi: 10.1146/annurev-environ-102014-021257.

Nilsson, R. H., Larsson, K. H., Taylor, A. F. S., Bengtsson-Palme, J., Jeppesen, T. S., Schigel, D., et al. (2019). The UNITE database for molecular identification of fungi: Handling dark taxa and parallel taxonomic classifications. Nucleic Acids Res. 47, D259–D264. doi: 10.1093/nar/gky1022.

op de Beeck, M., Lievens, B., Busschaert, P., Declerck, S., Vangronsveld, J., and Colpaert, J. v. (2014). Comparison and validation of some ITS primer pairs useful for fungal metabarcoding studies. PLoS ONE 9. doi: 10.1371/journal.pone.0097629.

Peham, T., Steiner, F. M., Schlick-Steiner, B. C., and Arthofer, W. (2017). Are we ready to detect nematode diversity by next generation sequencing? Ecol. Evol. 7, 4147–4151. doi: 10.1002/ece3.2998.

Penton, C. R., Gupta, V. V. S. R., Yu, J., and Tiedje, J. M. (2016). Size matters: Assessing optimum soil sample size for fungal and bacterial community structure analyses using high throughput sequencing of rRNA gene amplicons. Front. Microbiol. 7. doi: 10.3389/fmicb.2016.00824.

Quast, C., Pruesse, E., Yilmaz, P., Gerken, J., Schweer, T., Yarza, P., et al. (2013a). The SILVA ribosomal RNA gene database project: Improved data processing and web-based tools. Nucleic Acids Res. 41. doi: 10.1093/nar/gks1219.

Quast, C., Pruesse, E., Yilmaz, P., Gerken, J., Schweer, T., Yarza, P., et al. (2013b). The SILVA ribosomal RNA gene database project: Improved data processing and web-based tools. Nucleic Acids Res. 41. doi: 10.1093/nar/gks1219.

Quist, C. W., Gort, G., Mooijman, P., Brus, D. J., van den Elsen, S., Kostenko, O., et al. (2019). Spatial distribution of soil nematodes relates to soil organic matter and life strategy. Soil Biol. Biochem. 136. doi: 10.1016/j.soilbio.2019.107542.

R Core Team (2021). R: A language and environment for statistical computing. Available at: https://www.r-project.org/ [Accessed April 7, 2022].

Ranjard, L., Lejon, D. P. H., Mougel, C., Schehrer, L., Merdinoglu, D., and Chaussod, R. (2003). Sampling strategy in molecular microbial ecology: Influence of soil sample size on DNA fingerprinting analysis of fungal and bacterial communities. Environ. Microbiol. 5, 1111–1120. doi: 10.1046/j.1462-2920.2003.00521.x.

Reuter, J. A., Spacek, D. v., and Snyder, M. P. (2015). High-Throughput Sequencing Technologies. Mol. Cell 58, 586–597. doi: 10.1016/j.molcel.2015.05.004.

Sapkota, R., and Nicolaisen, M. (2015). High-throughput sequencing of nematode communities from total soil DNA extractions. BMC Ecol. 15. doi: 10.1186/s12898-014-0034-4.

Schoch, C. L., Seifert, K. A., Huhndorf, S., Robert, V., Spouge, J. L., Levesque, C. A., et al. (2012). Nuclear ribosomal internal transcribed spacer (ITS) region as a universal DNA barcode marker for Fungi. Proc Natl Acad Sci USA 109, 6241–6246. doi: 10.1073/pnas.1117018109.

Sikder, M. M., Vestergård, M., Sapkota, R., Kyndt, T., and Nicolaisen, M. (2020). Evaluation of metabarcoding primers for analysis of soil nematode communities. Diversity (Basel) 12, 1–14. doi: 10.3390/d12100388.

Song, Z., Schlatter, D., Kennedy, P., Kinkel, L. L., Kistler, H. C., Nguyen, N., et al. (2015). Effort versus reward Preparing samples for fungal community characterization in high-throughput sequencing surveys of soils. PLoS ONE 10. doi: 10.1371/journal.pone.0127234.

Stevens, J. L., Jackson, R. L., and Olson, J. B. (2013). Slowing PCR ramp speed reduces chimera formation from environmental samples. J. Microbiol. Methods 93, 203–205. doi: 10.1016/j.mimet.2013.03.013.

Štursová, M., Bárta, J., Šantrůčková, H., and Baldrian, P. (2016). Small-scale spatial heterogeneity of ecosystem properties, microbial community composition and microbial activities in a temperate mountain forest soil. FEMS Microbiol. Ecol. 92, fiw185. doi: 10.1093/femsec/fiw185.

Tanabe, A. S., and Toju, H. (2013). Two New Computational Methods for Universal DNA Barcoding: A Benchmark Using Barcode Sequences of Bacteria, Archaea, Animals, Fungi, and Land Plants. PLoS ONE 8. doi: 10.1371/journal.pone.0076910.

Tedersoo, L., Sánchez-Ramírez, S., Kõljalg, U., Bahram, M., Döring, M., Schigel, D., et al. (2018). High-level classification of the Fungi and a tool for evolutionary ecological analyses. Fungal Diversity 90, 135–159. doi: 10.1007/s13225-018-0401-0.

Thijs, S., de Beeck, M. O., Beckers, B., Truyens, S., Stevens, V., van Hamme, J. D., et al. (2017). Comparative evaluation of four bacteria-specific primer pairs for 16S rRNA gene surveys. Front. Microbiol. 8. doi: 10.3389/fmicb.2017.00494.

Thompson, L. R., Sanders, J. G., McDonald, D., Amir, A., Ladau, J., Locey, K. J., et al. (2017). A communal catalogue reveals Earth’s multiscale microbial diversity. Nature 551, 457–463. doi: 10.1038/nature24621.

Toju, H., Kurokawa, H., and Kenta, T. (2019). Factors Influencing Leaf- and Root-Associated Communities of Bacteria and Fungi Across 33 Plant Orders in a Grassland. Front. Microbiol. 10. doi: 10.3389/fmicb.2019.00241.

Toju, H., Tanabe, A. S., Yamamoto, S., and Sato, H. (2012). High-coverage ITS primers for the DNA-based identification of ascomycetes and basidiomycetes in environmental samples. PLoS ONE 7. doi: 10.1371/journal.pone.0040863.

Treonis, A. M., Unangst, S. K., Kepler, R. M., Buyer, J. S., Cavigelli, M. A., Mirsky, S. B., et al. (2018). Characterization of soil nematode communities in three cropping systems through morphological and DNA metabarcoding approaches. Sci. Rep. 8. doi: 10.1038/s41598-018-20366-5.

van den Hoogen, J., Geisen, S., Routh, D., Ferris, H., Traunspurger, W., Wardle, D. A., et al. (2019). Soil nematode abundance and functional group composition at a global scale. Nature 572, 194–198. doi: 10.1038/s41586-019-1418-6.

Waeyenberge, L., de Sutter, N., Viaene, N., and Haegeman, A. (2019). New insights into nematode DNA-metabarcoding as revealed by the characterization of artificial and spiked nematode communities. Diversity (Basel) 11. doi: 10.3390/d11040052.

Wall, D. H., Nielsen, U. N., and Six, J. (2015). Soil biodiversity and human health. Nature 528, 69–76. doi: 10.1038/nature15744.

Wiesel, L., Daniell, T. J., King, D., and Neilson, R. (2015). Determination of the optimal soil sample size to accurate19ecko19terizeise nematode communities in soil. Soil Biol. Biochem. 80, 89–91. doi: 10.1016/j.soilbio.2014.09.026.

Yeates, G. W., Wardle, D. A., and Watson, R. N. (1999). Responses of soil nematode populations, community structure, diversity and temporal variability to agricultural intensification over a seven-year period. Soil Biol. Biochem. 31, 1721–1733. doi: 10.1016/S0038-0717(99)00091-7.

Zhang, Y., Ji, L., and Yang, L. (2021). Abundance and diversity of soil nematode community at different altitudes in cold-temperate montane forests in northeast China. Glob. Ecol. Conserv. 29, e01717. doi: 10.1016/j.gecco.2021.e01717.

Zielińska, S., Radkowski, P., Blendowska, A., Ludwig-Gałęzowska, A., Łoś, J. M., and Łoś, M. (2017). The choice of the DNA extraction method may influence the outcome of the soil microbial community structure analysis. Microbiologyopen 6, e00453. doi: 10.1002/mbo3.453.

