## Supplemental Figure 1 for "Effects of source sample amount on biodiversity surveys of bacteria, fungi, and nematodes in soil ecosystems"

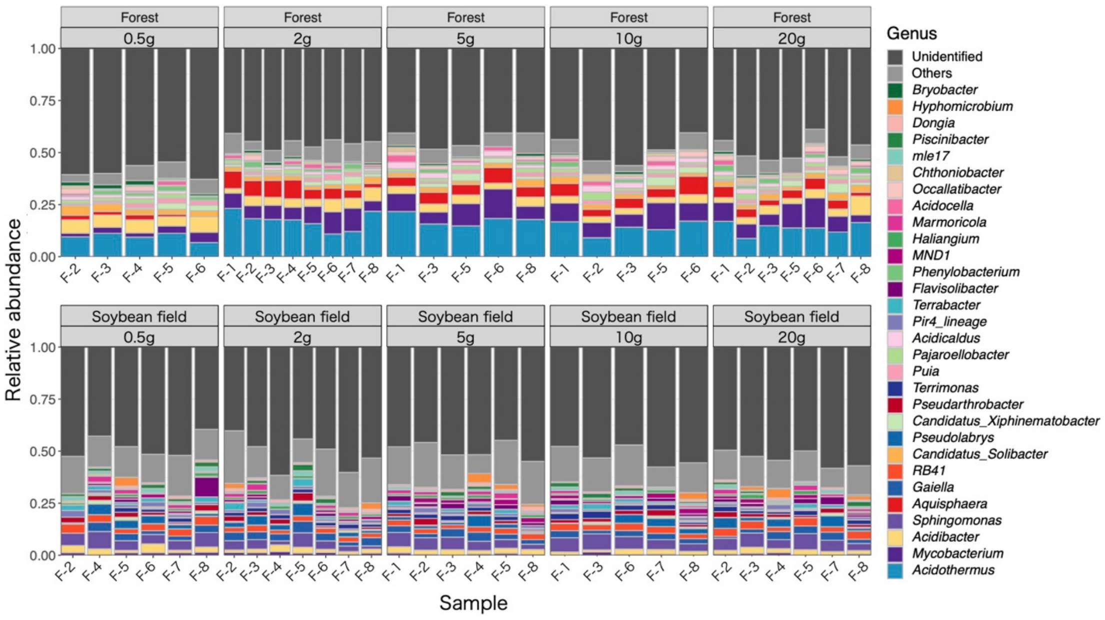


Supplementary Figure 1. Genus-level taxonomic compositions of bacteria. For each replicate sample of each ecosystem type (forest or soybean field), genus-level compositions of the sequencing data are shown at each soil-amount class. Samples with less than 1,000 sequencing reads were omitted.
