## Supplemental Figure 6 for "Effects of source sample amount on biodiversity surveys of bacteria, fungi, and nematodes in soil ecosystems"

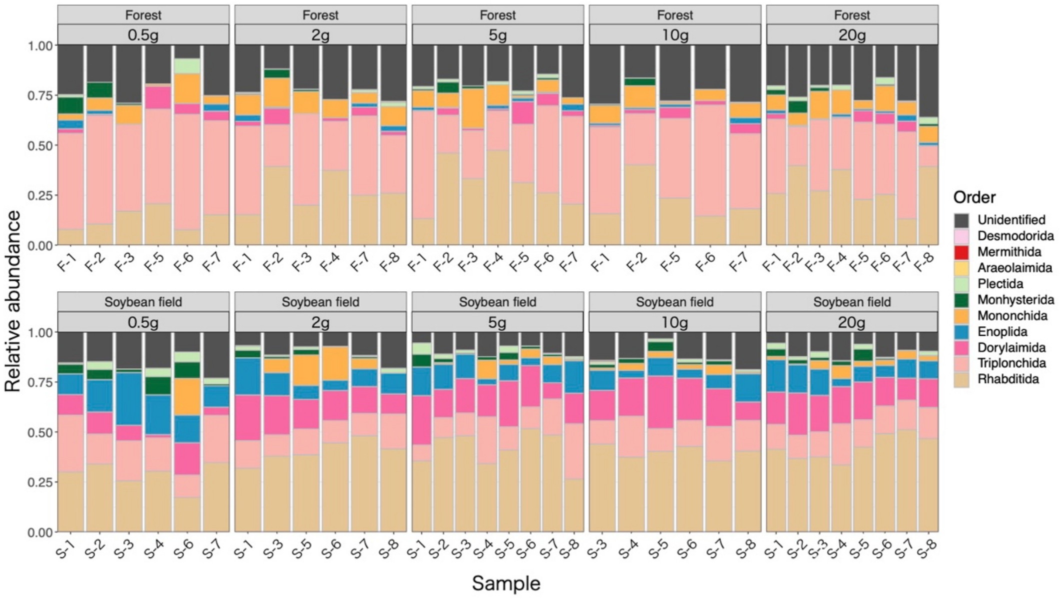


**Supplementary Figure 6.** Order-level taxonomic compositions of bacteria. For each replicate sample of each ecosystem type (forest or soybean field), order-level compositions of the sequencing data are shown at each soil-amount class. Samples with less than 1,000 sequencing reads were omitted.
