## Supplemental Figure 7 for "Effects of source sample amount on biodiversity surveys of bacteria, fungi, and nematodes in soil ecosystems"

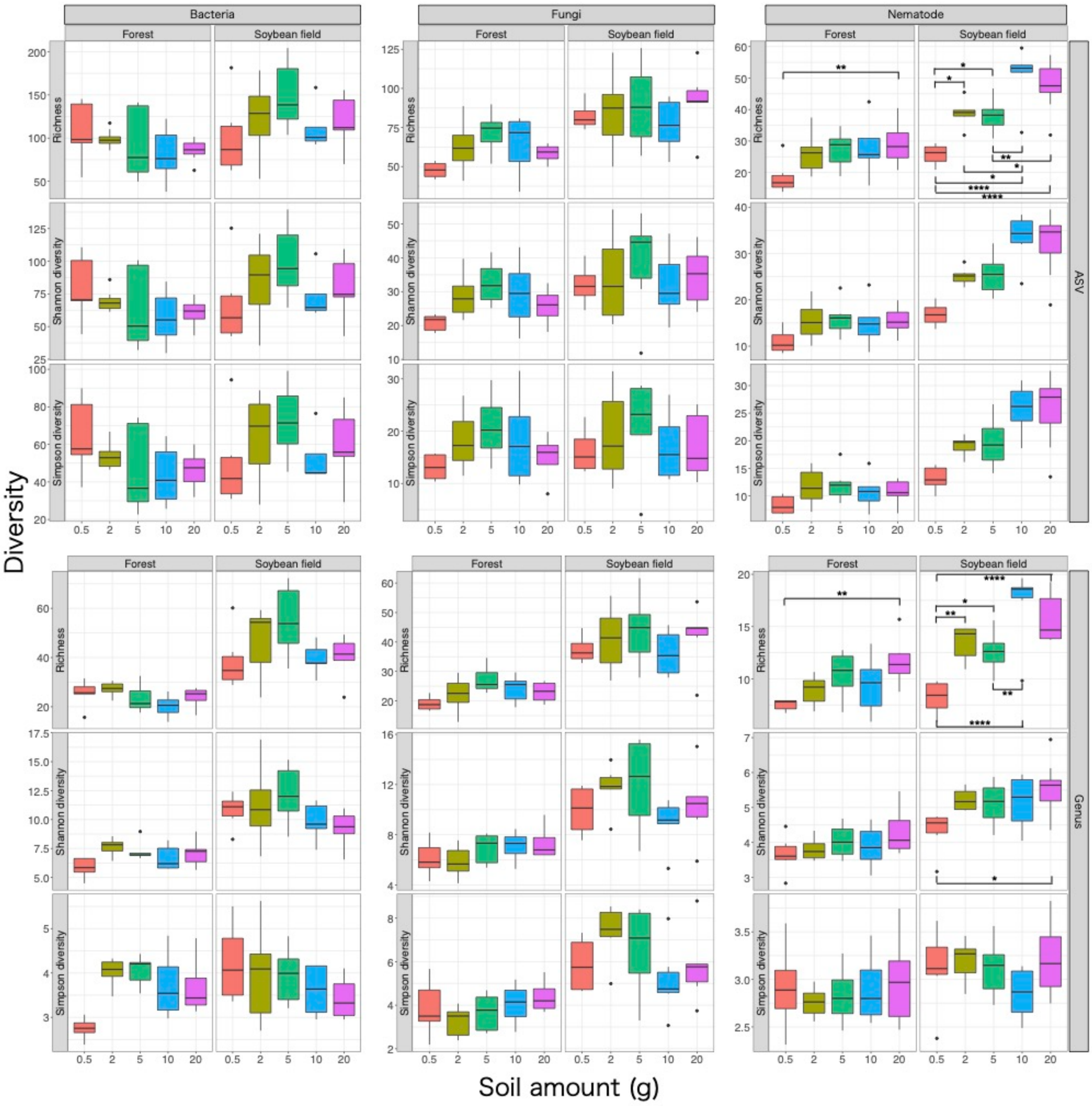


**Supplementary Figure 7.** Effects of soil amount in DNA extraction on ASV- and genus-level $\alpha$-diversity. For each of the bacterial 16S rRNA, fungal ITS, and nematode 18S rRNA datasets, taxonomic richness, Shannon diversity, and Simpson diversity at the ASV- and genus-level are shown with boxplots for ecosystem type (forest or soybean field).
